# A simple, cost-effective and automation-friendly direct PCR approach for bacterial community analysis

**DOI:** 10.1101/2021.03.01.433496

**Authors:** Fangchao Song, Jennifer V. Kuehl, Arjun Chandran, Adam P. Arkin

**Author notes:** (FS); (APA).

## Abstract

Bacterial communities in water, soil, and humans play an essential role in environmental ecology and human health. PCR-based amplicon analysis, such as 16S ribosomal RNA sequencing, is a fundamental tool for quantifying and studying microbial composition, dynamics, and interactions. However, given the complexity of microbial communities, a substantial number of samples becomes necessary to analyses that parse the factors that determine microbial composition. A common bottleneck in performing these kinds of experiments is genomic DNA (gDNA) extraction, which is time-consuming, expensive, and often biased on the types of species. Direct PCR methods are a potentially simpler and more accurate alternative to gDNA extraction methods that do not require the intervening purification step. In this study, we evaluated three variations of direct PCR methods using diverse heterogeneous bacterial cultures, ZymoBIOMICS Microbial Community Standards, and groundwater. By comparing direct PCR methods with DNeasy blood and tissue kits and DNeasy Powersoil kits, we found a specific variant of the direct PCR method exhibits a comparable overall efficiency to the conventional DNeasy Powersoil protocol. We also found the method showed higher efficiency for extracting gDNA from the gram negative strains compared to DNeasy blood and tissue protocol. This direct PCR method is 1600 times cheaper ($0.34 for 96 samples), 10 times simpler (15 min hands-on time for 96 samples) than DNeasy Powersoil protocol. The direct PCR method can also be fully automated, and is compatible with small volume samples, thereby permitting scaling of samples and replicates needed to support high-throughput large-scale bacterial community analysis.

**IMPORTANCE:** Understanding bacterial interaction and assembling in complex microbial communities using 16S ribosomal RNA sequencing normally requires a large experimental load. However, the current DNA extraction methods including cell disruption and genomic DNA purification are normally biased, costly, time and labor consuming, and not amenable to miniaturization by droplets or 1536 well plates due to the significant DNA loss during purification step for tiny volume and low cell density samples. Direct PCR method could potentially solve these problems. In this study, we demonstrate a direct PCR method which exhibits similar efficiency as the widely used method – DNeasy Powersoil protocol, while 1600 times cheaper and 10 times faster to execute. This simple, cost-effective, and automation friendly direct PCR based 16S ribosomal RNA sequencing method allows us to study the dynamics, microbial interaction and assembly of varying microbial communities in a high throughput fashion.

## Introduction

The microbial communities that populate water, soil, and animals drive complex ecological processes and play influential roles in ecosystem services and health. Studying these processes often starts by assessing bacterial diversity and identifying bacterial species. For complex communities this is most often accomplished by amplifying 16S ribosomal RNA from microbiome samples with PCR targeting specified variable regions, sequencing the complex mixture of molecules, and calculating the relative abundance of the inferred distinct ribosome genes (16S sequencing).^1–3^ It is often the goal to use these data to find associations between community composition and biological function. However, this is hampered by a number of complexities in analysis and interpretation of these data arising at multiple stages of the process. These can range from biased extraction and amplification of nucleic acids from different types of bacteria in different growth phases to problems of abundance estimation. Amongst them, DNA extraction is considered to lead to the most striking bias between previously tested protocols.^4, 5^ Moreover, research on microbiomes is rapidly expanding, while the cost of such research, due to the widespread use of laboratory automation and the progress of next generation sequencing, is decreasing. Large scale microbial community analysis is hampered by the time and labor cost of DNA extraction and purification.^6^ In addition, as high throughput bacterial assays get smaller, such as in droplet or in well plate based assays, current column or beads based DNA purification method are ineffective due to significant DNA loss for low cell number samples.^7–10^ Therefore, it is imperative to design a new DNA extraction and amplification method which is efficient, cost-effective, and amenable to miniaturization.

There have been a number of DNA extraction approaches to increase the general efficiency of extracting nucleic acids (DNA and RNA) from all cells in a sample.^11–14^ These methods differ based on organism type, sample materials (e.g. sediment vs. water), and the compatibility for downstream processing. Ideally, cell disruption and DNA amplification could be done without an intervening purification step, and the entire set of operations should be amenable to miniaturization and automation. However, these “direct PCR” techniques, while popular, when applied to bacteria are thought to be non-quantitative, and worse than the “non-direct PCR” methods, due to the limited choices of bacterial cell disruption methods compatible with PCR chemistries. Of those that have become popular, very few have been rigorously tested both for efficiency and precision in the application of 16S sequencing of microbial communities.^15, 16^ None of the direct PCR methods have been quantified by comparing the obtained composition with the real composition (of a standard community) and have not been optimized for automation or miniaturization. Table 1 lists some of the basal cell disruption methods used for DNA extraction along with their functions and PCR compatibility. Amongst them, alkaline and beads-beating are very effective universal disruptors but DNA released early during these processes may be damaged by the duration of treatments necessary to extract DNA from more recalcitrant cells.^17^ Sodium dodecyl sulfate (SDS) is very effective for gram negative bacteria but its use generally requires a purification step before PCR amplification of the sample.^18^ Only a small portion of these methods are compatible with direct PCR, of which very few have been directly tested on samples of known composition with sufficient diversity to uncover biases in extraction. Thus we propose that the combination of a few of the basal DNA disruption methods would fulfill the needs of an effective direct PCR protocol for extraction and amplification of 16S RNA from bacterial microbiome samples.

**Table 1.** List of cell disruption methods, functions, and PCR compatibility.

| Cell lysis methods |  | Functions | PCR compatible or not | References |
| --- | --- | --- | --- | --- |
| Mechanical | Beads beating | mechanically destroy cell membrane structure | Yes | 34,35 |
|  | Freeze-thaw | Cell membrane cracking by ice crystals | Yes | 34,36 |
| Chemical | Sodium dodecyl sulfate (SDS), anionic surfactant | Disrupt the cell membrane phospholipids, strong | Generally no, conditionally yes | 34,37 |
|  | Sodium lauroyl sarcosinate (Sarkosyl), anionic surfactant | Disrupt the cell membrane phospholipids, strong | No | 38,39 |
|  | Cetyl trimethylammonium bromide (CTAB), quaternary ammonium surfactant | Disrupt the cell membrane phospholipids, strong | Unknown | 34,40 |
|  | IGEPAL CA-630, non-ionic surfactant | Disrupt the cell membrane phospholipids, mild | Yes | 19,20 |
|  | Triton X-100, non-ionic surfactant | Disrupt the cell membrane phospholipids, mild | Yes | 34,41 |
|  | CHAPS (3-cholamidopropyl dimethylammonio 1-propanesulfonate), zwitterionic surfactant | Disrupt cell membrane | No | 41,42 |
|  | Potassium hydroxide, alkaline | Disrupt cell membrane, strong | No | 43,44 |
| Enzymatic | Lysozyme | Disrupt Peptidoglycan | No | 34,45 |
|  | Lysostaphin | Disrupt Peptidoglycan | Unknown | 46,47 |
|  | Proteinase K | Disrupt membrane protein | Yes by deactivation | 18,34 |
| Heat | Boiling | Disrupt cell membrane and protein denaturation | Yes | 48,49 |

We sought an effective combination of DNA extraction techniques that were cheap, easy and compatible with direct PCR to reduce bias and increase scalability. To do so, we explored the use of a PCR-compatible non-ionic surfactant (IGEPAL CA-630) that has been successfully applied in eukaryotic proteomics and RNA-SEQ studies,^19, 20^ with other PCR-compatible techniques such as freeze-thaw cycles, proteinase K treatment, and variation in heating time. We compare the performance of IGEPAL treatments with different combinations of these three membrane disruption methods to each other, and two commerical kits-the DNeasy Blood and Tissue kit and the DNeasy Powersoil DNA isolation kit (Powersoil) which are two of the most widely used methods for effective DNA extraction. To assess performance, we test the methods on a mock community designed to encompass bacteria with different cellular properties in different phases of growth and on a more diverse groundwater community. We found that the best combination of our approach yields overall comparable quantification to Powersoil method, though biases still persist (like Powersoil method), but with a far shorter and more cost-effective protocol that is more compatible with miniaturization and automation.

## Results

We evaluated the three new protocols shown in Figure 1: The IGEPAL only method (Method 1) uses the surfactant (IGEPAL CA-630) and longer heating than current protocols (10 min at 98°C for initial activation during PCR); the IGEPAL+Freeze-thaw method (Method 2) adds freeze-thaw sequence on top of Method 1; IGEPAL+Freeze-thaw+ProteinaseK method (Method 3) adds a proteinase K treatment and extra heating during proteinase K treatment on top of Method 2. Since the three protocols use different combinations of mechanisms to disrupt membranes, we expect increasing extraction efficiency and decreasing bias as we go from Method 1 to 3. As shown in Table 1, surfactant may disrupt the membrane phospholipids; proteinase K disrupts membrane proteins; heating and freeze-thaw generally disrupt the bacterial membrane mechanically. To evaluate these methods, we first quantify the extraction efficiency on a set of specially chosen target bacteria with different membrane properties using quantitative PCR (qPCR). We then compare these methods to Powersoil kit by quantifying the ability to operate in mixed culture and reproduce known abundance ratios using a specially designed mock community. We further compare the results from application to diverse groundwater communities sampled from the Bear Creek Valley watershed of Oak Ridge, TN.. Finally, the cost and other aspects of these methods are evaluated in Table 2.

**Table 2.** Comparison of the DNA extraction and direct PCR methods.

| Method | Cost/96 samples (USD)<sup>a</sup> | Protocol steps<sup>b</sup> | Extraction time<sup>c</sup> | Hands-on time<sup>c</sup> |
| --- | --- | --- | --- | --- |
| DNeasy Powersoil HTP 96 kit | \$552.7 | 32 | 4 h | 3.5 h |
| Extract-N-Amp Plant PCR kit | \$176.9 | 3 | 25 min | 15 min |
| Method 1 (IGEPAL only) | \$0.004 | None | Negligible | Negligible |
| Method 2 (IGEPAL+Freeze-thaw) | \$0.012 | 2 | 45 min | 10 min |
| Method 3 (IGEPAL+Freeze-thaw+ProteinaseK) | \$0.34 | 3 | 2 h | 15 min |

<sup>a</sup>The calculation of the cost refer to Table S3. <sup>b</sup>If there is no extra step before adding PCR reagents, it is none. <sup>c</sup>Time is estimated for processing 96 samples including waiting time, the actual labor may be different. Please check the discussion in the main text.

**Figure 1.**
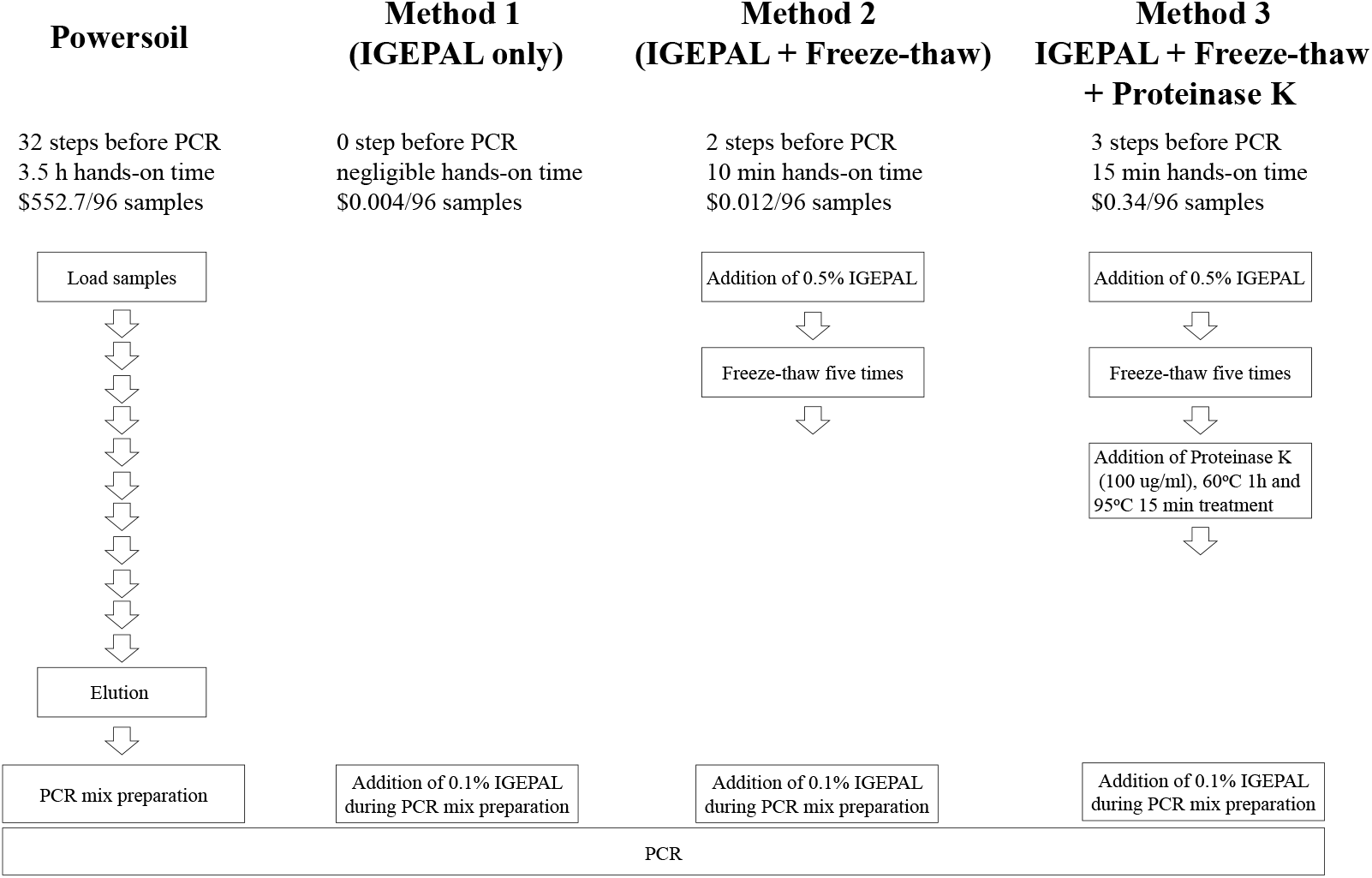
Workflows of conventional DNA extraction method and direct PCR methods for analyzing microbial community.

### Evaluation of direct PCR methods using model strains and quantitative PCR (qPCR)

The structure of microbial membranes and other elements that prevent DNA accessibility are extremely diverse among bacteria phyla, and can even vary across the growth phases of a given species.^21, 22^ Therefore, it is almost inevitable that any method of DNA extraction will have a different efficacy across these factors. To understand the variation of efficacy, we start by testing our methods on four model strains - two gram negative strains (*Escherichia coli* and *Pseudomonas putida*) and two gram positive strains (*Lactococcus lactis subsp.* and *Lactobacillus brevis*) in both exponential and stationary phase.

We first tested if IGEPAL alone could efficiently kill bacteria and be compatible with PCR. Stationary phase cultures of our test bacteria (*Escherichia coli*, *Pseudomonas putida*, and *Lactococcus lactis subsp.*) were exposed to 0.1% IGEPAL at 98°C for 5 min and were plated on LB agar plates. No colonies appeared after two days growth implying full efficacy in killing the bacteria. We then tested PCR of three different 16S PCR primers using a standard protocol with either KAPA HotStart HiFi polymerase or Taq polymerase augmented with 0.1% IGEPAL. The gel image of PCR products shown in Figure S1(A) indicates that the PCR was unperturbed by addition of the surfactant - IGEPAL CA-630.

We then evaluated the ability of the three direct PCR methods to quantify the amount of gDNA in each exponential and stationary phase sample compared to one of the most popular commercial kits - the DNeasy Blood and Tissue kit which has been recommended by comparing with multiple commercial available kits, for complete gDNA extraction.^11, 23^ We compare the estimation of relative gDNA concentrations using the threshold cycle (Ct) of qPCR by the direct PCR methods vs the DNeasy Blood and Tissue kit using the same cell cultures. Figure 2 shows the direct comparison of Ct between gDNA extracted by using the DNeasy Blood and Tissue kit (gDNA Control) and an equivalent amount of cells with the direct PCR methods. To better compare the difference between the gDNA control and the direct PCR methods, the average difference of Ct between direct PCR methods and gDNA control (ΔCt = Ct of direct PCR - Ct of gDNA control) and the *p* value by *t* test are calculated and shown in Table S1. In Figure 2, the cell types with ΔCt > 0 and *p* < 0.05 are labeled with “+”, which represents the direct PCR methods are less effective compared to the gDNA control; the cell types with ΔCt < 0 and *p* < 0.05 are labeled with “−”, which indicates the direct PCR methods are more effective compared to the DNeasy Blood and Tissue kit; the cell types with *p* > 0.05 are not labeled, which suggests the direct PCR methods seem to be similarly effective compared to the DNeasy Blood and Tissue kit.

**Figure 2.**
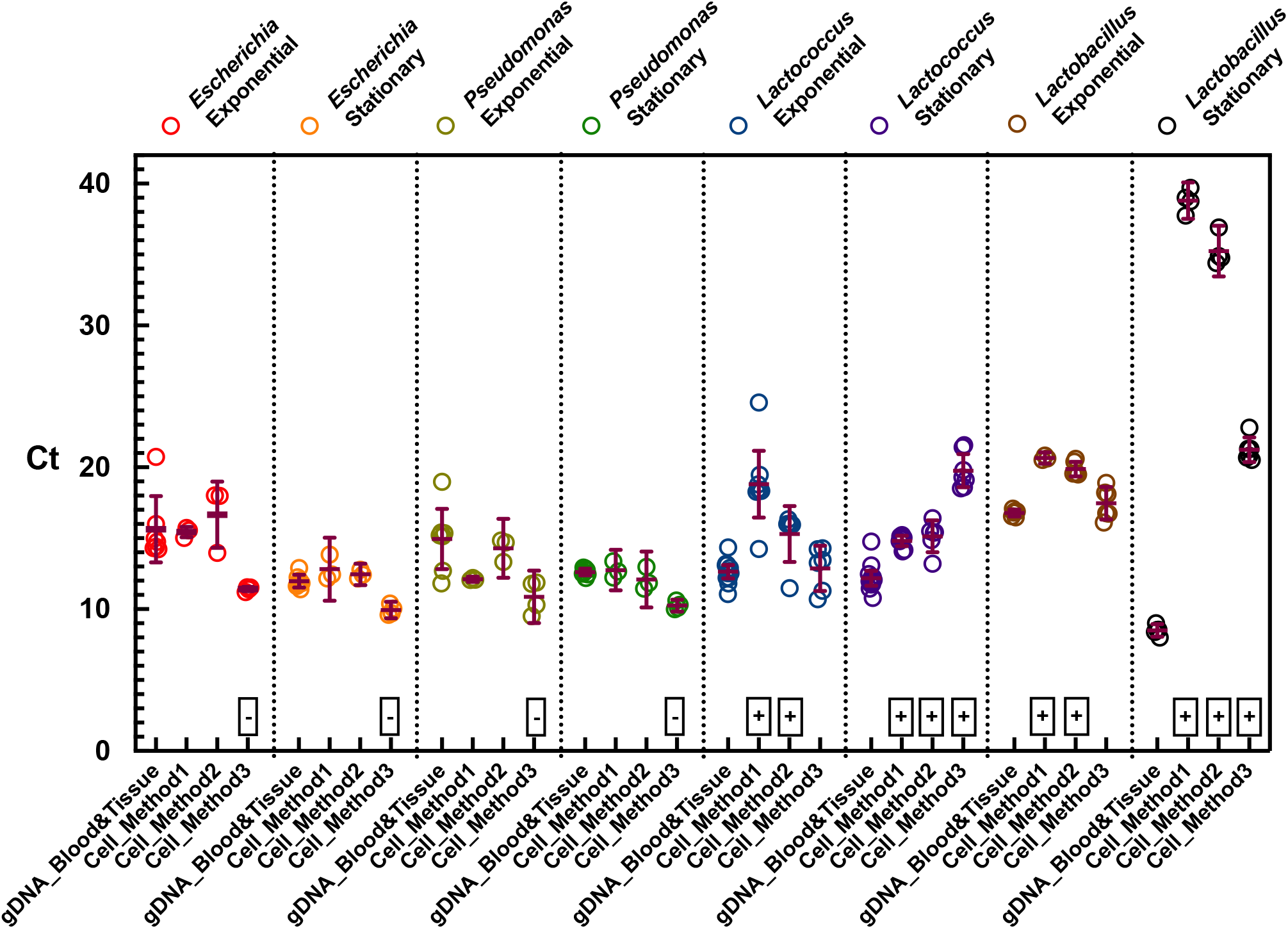
Ct from quantitative PCR (qPCR) of genomic DNA extracted by DNeasy blood and tissue kits and cells treated by direct PCR methods (Method 1: IGEPAL only; Method 2: IGEPAL + Freeze-thaw; Method 3: IGEPAL + Freeze-thaw + proteinase K). The strains are *Escherichia coli str. K-12 substr.* MG1655 (*E. coli*), *Pseudomonas putida* KT2440 (*P. putida*), *Lactococcus lactis subsp. cremoris str.* MG1363 (*L lactis*), and *Lactobacillus brevis* ATCC 14869 (*L. brevis*). + represents Ct of the direct PCR methods > Ct of the extracted gDNA; - represents Ct of the direct PCR methods < Ct of the extracted gDNA; others indicate Ct are similar between extracted gDNA and direct PCR methods.

Collectively, all the three methods could effectively interrupt the gram negative bacteria (*Escherichia coli*, *Pseudomonas putida*) in both exponential and stationary phases, similar to or better than the DNeasy Blood and Tissue kit. In particular, *Escherichia coli* and *Pseudomonas putida* in both exponential and stationary phases lysed by Method 3 exhibit smaller Ct than the gDNA control. It indicates that more gDNA is extracted by Method 3 than the DNeasy Blood and Tissue kit from the same amount of cell culture. Thus, Method 3 (IGEPAL+Freeze-thaw+ProteinaseK) exhibits better results compared to DNeasy Blood and Tissue kit for gram negative strains. This is probably because the direct PCR method avoids gDNA loss due to binding on the spin-column of the DNeasy Blood and Tissue kit. Methods 1 (IGEPAL only) and 2 (IGEPAL+Freeze-thaw method) exhibit similar efficiency for extracting gDNA from gram negative bacteria (*Escherichia coli*, *Pseudomonas putida*) in both exponential and stationary phases.

The gDNA of exponential phase cultures of *Lactococcus lactis subsp.* and *Lactobacillus brevis* is as effectively extracted by Method 3 (IGEPAL+Freeze-thaw+ProteinaseK), as from the gDNA control (DNeasy Blood and Tissue kit). As shown in Figure 2 and Table S1, the ΔCt of exponential phase cultures of *Lactococcus lactis subsp.* (ΔCt Δ 0.19 *±* 0.7 and *p* = 0.787) and *Lactobacillus brevis* (ΔCt = 0.70 *±* 0.5 and *p* = 0.19) extracted by Method 3 is close to their gDNA control extracted by DNeasy Blood and Tissue kit. Both Method 1 and Method 2 are less efficient in extracting gDNA of exponential phase cultures of *Lactococcus lactis subsp.* and *Lactobacillus brevis* than DNeasy Blood and Tissue kit. For examples, gDNA of exponential phase cultures of *Lactococcus lactis subsp.* and *Lactobacillus brevis* extracted by Method 2 is 4.6 (ΔCt = −2.56 *±* 0.86, *p* = 0.016) and 7.6 (ΔCt = −3.11 *±* 0.24, *p* < 0.0001) fold lower than DNeasy Blood and Tissue kit, respectively. In addition, the direct PCR methods have difficulties with effectively extracting gDNA from the stationary phase cultures of both *Lactococcus lactis subsp.* and *Lactobacillus brevis*, in particular for method 1 (ΔCt = −3.11 *±* 0.24 for *Lactococcus lactis subsp.* and ΔCt = −3.11 *±* 0.24 for *Lactobacillus brevis* with *p* < 0.0001 for both.). Apparently, the surfactant could not disrupt the thick peptidoglycan in the cell wall of gram positive bacteria. Method 3 improves the efficiency significantly by adding both freeze-thaw and proteinase K treatment. As expected, the efficiency of cell disruption increases from method 1 to 3. In Figure S1(B), the qPCR curves of the exponential phase *Lactococcus lactis subsp.* from Method 1 to 3 shift from right to left, getting close to the gDNA control. And the cell cultures of exponential phase *Lactococcus lactis subsp.* treated by the direct PCR method from Method 1 to 3 turn to be more transparent as shown in Figure S1(C). To further improve the gDNA extraction of stationary phase gram positive bacteria, lysozyme treatment - 10 mg/ml lysozyme treatment at 55oC for 20 min was applied before proteinase K treatment in Method 3. In this design, the lysozyme disrupts the peptidoglycan, and proteinase K denatured lysozyme to eliminate the potential inhibitive effects on PCR. However, either lysozyme or proteinase K-denatured lysozyme inhibits PCR (gel image is not shown.).

### Evaluation of direct PCR methods using mock microbial community standard

To evaluate the ability of the direct PCR methods to accurately estimate the relative abundance of members in more complex microbial communities, we tested them against the most widely used microbiome gDNA extraction kit - the DNeasy Powersoil kit using a mock community standard - the ZymoBIOMICS Microbial Community Standard. The ZymoBIOMICS Microbial Community Standard is composed of three gram negative and five gram positive bacteria along with two yeast strains (not measured in our study). They differ in GC content, genome size, 16S copy number, and are mixed in a known ratio. The composition of ZymoBIOMICS Microbial Community Standard was measured by 16S sequencing either following gDNA extraction by Powersoil and 16S PCR, or following direct 16S PCR methods. The relative abundances of the standard community, and those measured by Powersoil and the three direct PCR methods are listed in Table S2.

All methods were able to extract DNA from all microbial species. The average composition of each species of the standard community, Powersoil, and the direct PCR methods are visualized in Figure 3(A). At a glance, the relative abundance obtained from Method 3 and Powersoil are much closer to the real composition of the standard community than those from Method 1 and 2. To quantify the similarity of the relative abundances obtained by the direct PCR methods and Powersoil to the real relative abundance of the standard community, we calculate the Euclidean distances between the relative abundance of each measurements and the real relative abundance of the standard community, and consider the Euclidean distances as the index of dissimilarity. In Figure 3(B), the dissimilarity of the composition from Method 3 to the composition of the standard community (0.29 *±* 0.01) is close to the dissimilarity of the composition from Powersoil to the composition of the standard community (0.22 *±* 0.05), although it is not exactly the same (*p* = 0.02, Welch’s *t* test). The dissimilarity between the composition from Method 1 and the composition of the standard community and the dissimilarity between the composition from Method 2 and the composition of the standard community is 0.52 *±* 0.04 and 0.42 *±* 0.01, respectively. Both Method 1 and Method 2 have a much higher bias compared to either Method 3 (0.29 *±* 0.01) and Powersoil (0.22 *±* 0.05). Figure 3(C) is the PCA plot of the relative abundances obtained from different methods compared to the real relative abundance of the standard community, in which the same symbols represent the replicates of each methods. Principle component 1 (PC1) which takes 83% weight dominates the differences.

**Figure 3.**
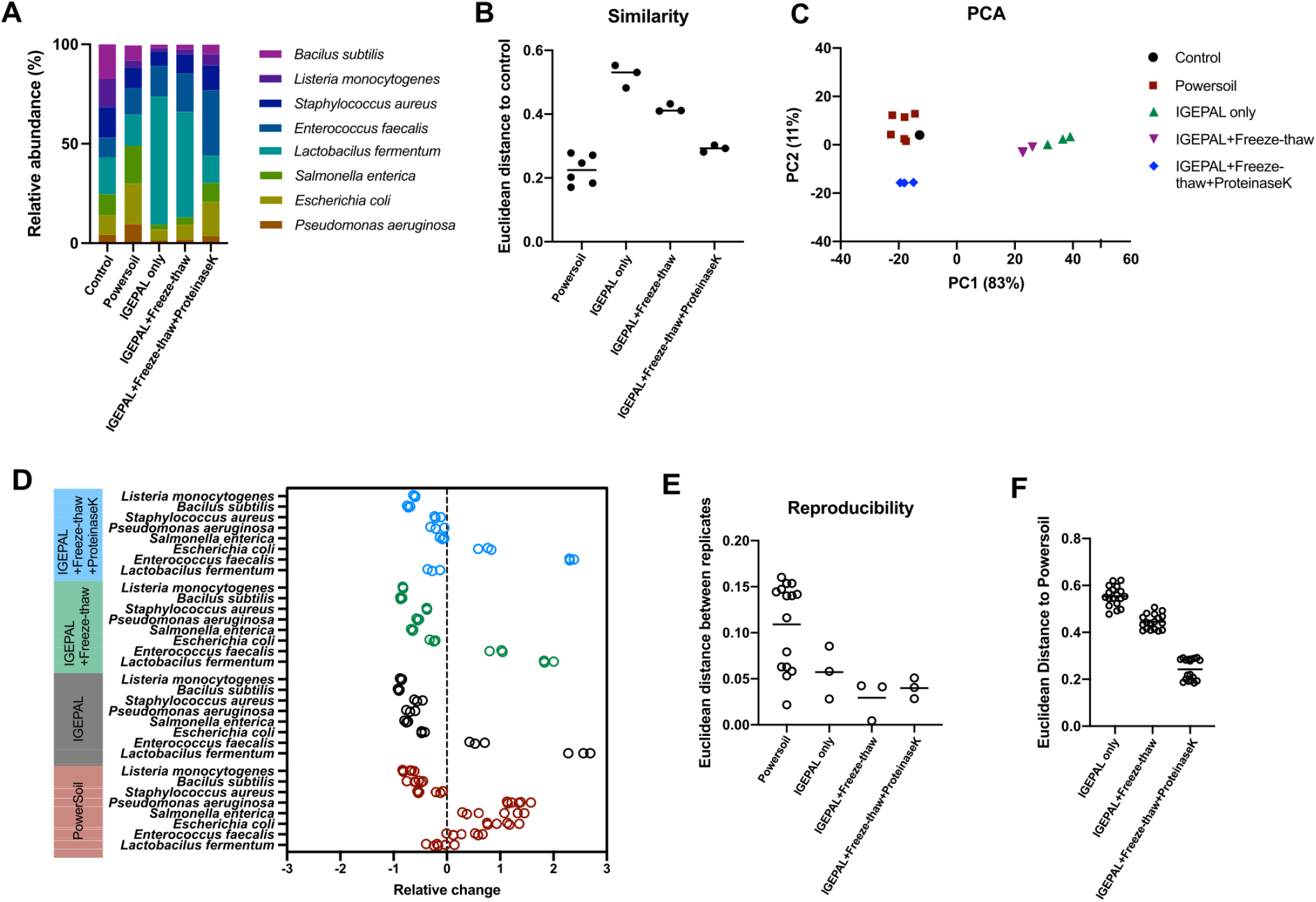
Evaluation of direct PCR methods and DNeasy Powersoil kits using ZymoBIOMICS Microbial Community Standards. Notes: IGEPAL only is Method 1; IGEPAL + Freeze-thaw is Method 2; IGEPAL + Freeze-thaw + proteinase K is Method 3. (A) Bacterial average composition of ZymoBIOMICS microbial community standards and those obtained by Powersoil and direct PCR methods. (B) The Euclidean distance of the relative abundance obtained from direct PCR methods and Powersoil to the real relative abundance of the standard community. (similarity to the real composition) (C) PCA plot derived from Euclidean distances among the real relative abundance of the standard community, the relative abundance obtained from direct PCR methods and Powersoil. (D) The relative changes of the relative abundance of each bacteria in ZymoBIOMICS microbial community standards using direct PCR methods and Powersoil compared to the real relative abundance of the standard community. Positive changes indicate higher relative abundance in the methods compared to the real relative abundance of the standard community, and negative changes indicate lower relative abundance in the methods compared to the real relative abundance of the standard community. (E) The Euclidean distances among replicates. (repeatability of each method) (F) The Euclidean distance of the direct PCR methods to Powersoil. (similarity to Powersoil)

To further understand the bias on different bacteria among different methods, the relative changes of the relative abundance of each member by different methods compared to the real relative abundance of the standard community are shown in Figure 3(D). These relative changes of relative abundance compared to the real relative abundance of the standard community are defined as the fold bias 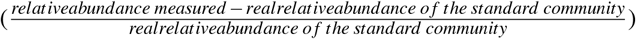. Almost all the bacteria showed less than one fold bias in Method 3 compared to the real relative abundance of the standard community, except *E. faecalis* which has 2.3 folds over-estimation of their relative abundance. In particular, the relative abundance bias of *S. aureus*, *P. aeruginosa*, *S. enterica*, and *L. fermentus* measured by Method 3 is very small: −0.18 *±* 0.07, −0.18 *±* 0.13, −0.09 *±* 0.04, and −0.25 *±* 0.12 folds of the relative abundance of the standard community, respectively. The relative abundances of three gram negative bacteria - *E. coli*, *P. aeruginosa*, and *S. enterica* measured by Powersoil have higher than one fold positive bias compared to the real relative abundance of the standard community. This demonstrates the systematic bias of gram negative preference of Powersoil DNA extraction kit. The relative changes of relative abundance of almost all the bacteria decreases from Method 1 (IGEPAL only) to Method 3 (IGEPAL+Freeze-thaw+ProteinaseK), except *E. faecalis*. It suggests that the gDNA extraction efficiency on most of the bacteria improves from Method 1 to 3, and *E. faecalis* seems to be the outlier. In addition, the gram positive bacteria *L. fermentus* with smallest genome size (1.905 Mb) in the mock community standard has the highest positive bias of relative abundance measured in Method 1 compared to the real relative abundance of the standard community, which is much higher than all the gram negative bacteria - *P. aeruginosa*, *E. coli*, and *S. enterica*. This indicates IGEPAL may be effective enough to disrupt some of the gram positive strains, like *L. fermentus*.

To determine the quantitative repeatability/reproducibility of replicate samples with the direct PCR methods and Powersoil, we use the Euclidean distances of the composition of each replicate measurement for each method (Figure 3(E)). Smaller euclidean distance indicates higher repeatability. And the direct PCR method 2 and 3 are more reproducible than Powersoil, since the euclidean distance score of Method 2 and Method 3 (0.057 *±* 0.028 and 0.029 *±* 0.022, respectively) are much lower than Powersoil (*p* = 0.003 for Powersoil compared to Method 2, *p* = 0.0002 for Powersoil compared to Method 3, Welch’s *t* test).

Since Powersoil is widely used in the microbiome gDNA extraction, we also compared the dissimilarity of the relative abundance measured by the direct PCR methods (Method 1, 2, and 3) to Powersoil using Euclidean distance as the index. As shown in Figure 3(F), the dissimilarity, defined as the Euclidean distance, to Powersoil decreases from Method 1 to 3 (*p* < 0.001, One way ANOVA). And the dissimilarity of the relative abundance obtained from Method 3 to Powersoil is 0.24 *±* 0.045, similar to the dissimilarity of the relative abundance from Powersoil to the real relative abundance of the standard community (0.22 *±* 0.05, *p* = 0.45, Welch’s *t* test).

### Evaluation of direct PCR methods by analyzing the microbiome in ground waters

To determine the practical outcomes that stem from differences in the above methods of quantifying microbial communities of environmental samples, we tested the direct PCR methods on ground water samples (GW822D and GW823E) from Bear Creek Valley watershed of Oak Ridge, TN. Since the “real” compositions of the communities of GW822D and GW823E are unknown, we evaluate the microbial composition obtained by the direct PCR methods compared to the microbial composition obtained by Powersoil which has been widely used and adopted as the standard protocol by The Earth Microbiome Project.^2, 15^ The relative abundances of exact sequence variants (ESVs) obtained by the direct PCR methods (Method 1, 2, and 3) and Powersoil are listed in Figure S2. The average composition of ESVs of Powersoil, and the direct PCR methods are visualized in Figure 4(A, B). The average composition in Phylum level is shown in Figure S3. At a glance, the composition obtained from direct PCR methods are all very similar to Powersoil. The dissimilarity is further quantified by the Bray-Curtis distance of each direct PCR method (1, 2, and 3) to Powersoil using the normalized and log transformed ESV metrics. The results are shown in Figure 4(C, D). The Bray-Curtis distance of the microbial composition of GW822D obtained from Method 1, 2, and 3 to Powersoil is 0.12 *±* 0.023, 0.076 *±* 0.028, and 0.12 *±* 0.010, respectively. For sample GW823E, the Bray-Curtis distance obtained from Method 1, 2, and 3 to Powersoil is 0.20 *±* 0.052, 0.096 *±* 0.031, and 0.11 *±* 0.033, respectively. The distance between Method 1 and Powersoil is higher than the distances between Method 2 or 3 and Powersoil (*p* < 0.05 for both, Welch’s *t* test). The principal-coordinate analysis (PCoA) based on the Bray-Curtis distances is shown in Figure 4(E). The dissimilarity is highly depended on the principle component 1 (87%). The data is strongly clustered according to the sample types (GW822D and GW823E), with minor separations among different methods. All of the direct PCR methods are able to reflect the differences of these two water samples (PERMANOVA, *p* < 0.001 by comparing samples). However, Method 2 and 3 are closer to the results obtained from the Powersoil method. We also evaluated the differences in alpha diversity (Shannon index) of samples GW822D and GW823E between the direct PCR and Powersoil method, which is shown in Figure 4(F). The alpha diversities are different between samples GW822D and GW823E (*p* = 0.002, Welch’s *t* test), but there are no significant differences across different methods (*p* > 0.3 for both GW822D and GW823E, one-way ANOVA).

**Figure 4.**
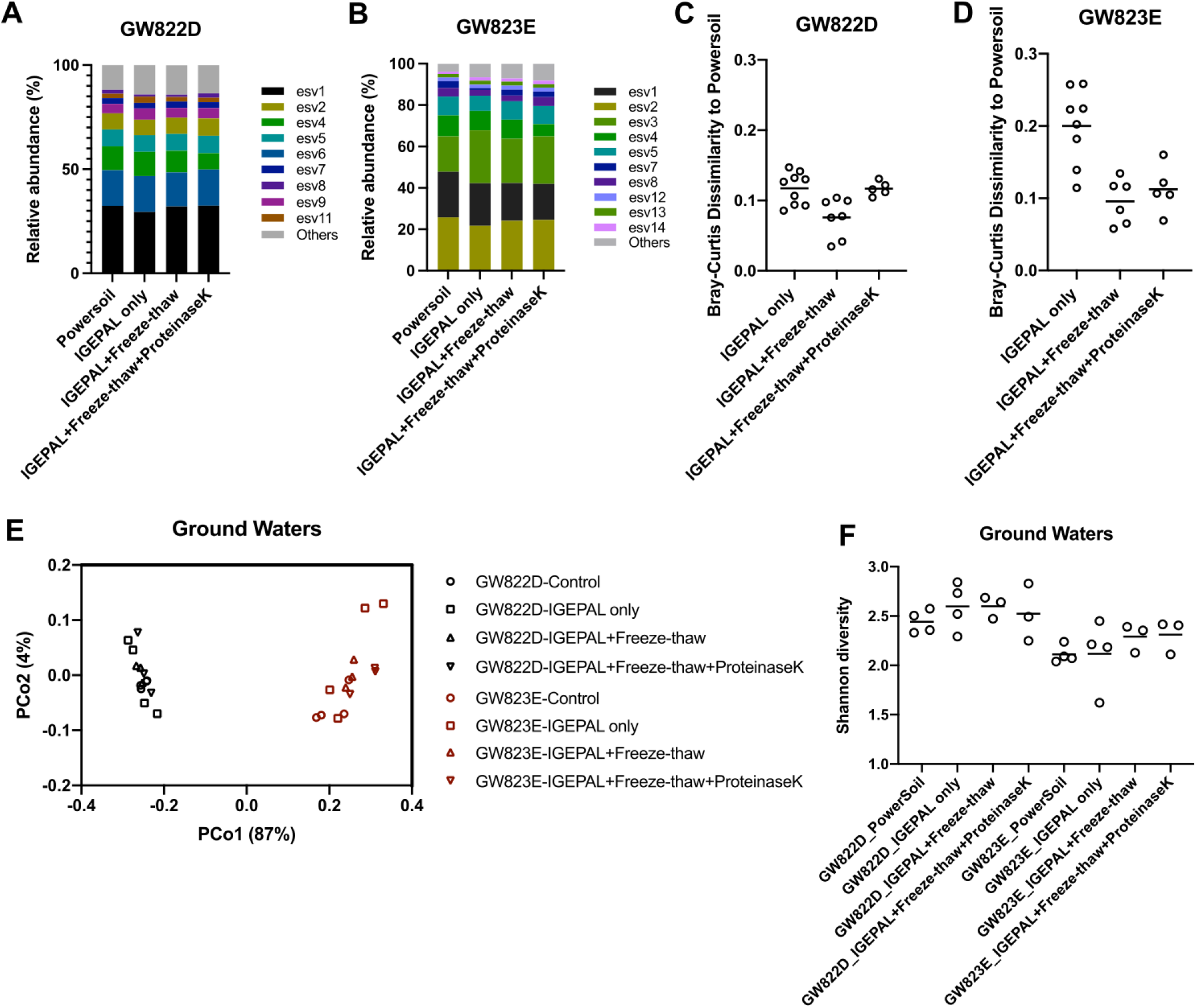
Comparison of direct PCR methods to Powersoil kits using ground water samples. Notes: IGEPAL only is Method 1; IGEPAL + Freeze-thaw is Method 2; IGEPAL + Freeze-thaw + proteinase K is Method 3. (A) Average bacterial Exact Sequence Variants (ESV) composition of the ground water sample GW822D. (B) Average bacterial ESV composition of the ground water sample GW823E. (C) The Bray-Curtis distances of the composition of GW822D obtained by direct PCR methods to Powersoil. (D) The Bray-Curtis distances of the composition of GW823E obtained by direct PCR methods to Powersoil. (E) PCoA plots derived from Bray-Curtis dissimilarities of the community compositions obtained using the direct PCR methods and Powersoil. (F) Alpha diversity (Shannon index) of GW822D and GW823E obtained by Powersoil and direct PCR methods.

### Some aspects of the new direct PCR methods

The efficiency and repeatability of the direct PCR methods have been evaluated via both testing on the mock microbial community standard, and comparing the direct PCR methods against DNeasy Blood and Tissue DNA extraction kit and DNeasy Powersoil DNA extraction kit. Like all the DNA extraction methods, the direct PCR methods have biases. However, the efficiency of the direct PCR methods, in particular Method 3, are comparable with the widely used DNA extraction kits - DNeasy Powersoil kit and DNeasy Blood and Tissue kit.

The direct PCR methods are more cost-effective, simple, and automation friendly. Table 2 summarizes the cost, complexity, and compatibility of DNeasy Powersoil kit, the direct PCR methods (Method 1, 2, and 3), and the Extract-N-Amp Plant PCR kit which was a previous reported direct PCR method having comparable efficiency with Powersoil as well.^15, 16^ The cost of the direct PCR methods, which are $0.004/96 samples for Method 1, $0.012/96 samples for Method 2, and $0.34/96 samples for Method 3, are all negligible and much lower than the cost of DNeasy Powersoil HTP 96 kit ($552.7/96 samples) and Extract-N-Amp Plant PCR kit ($176.9/96 samples). The cost analysis in detail is listed in Table S3. In particular, as the best performing direct PCR method, Method 3 (IGEPAL+Freeze-thaw+ProteinaseK) is 1600 times cheaper ($0.34/96 samples) than Powersoil ($552.7/96 samples), and 500 times cheaper than the cost of Extract-N-Amp ($176.9/96 samples).

The protocol of direct PCR is also simpler and shorter than Powersoil (Figure 1). Method 1 does not require any extra steps except adding IGEPAL in the PCR reagent. Method 2 contains the addition of IGEPAL and a freeze-thaw step. Method 3 adds proteinase K treatment on top of Method 2. And Extract-N-Amp Plant PCR kit has extraction solution addition, heating treatment, and dilution solution addition steps.^15^ Videvall et al. also have an additional shake step using TissueLyzer (Qiagen) on top of the Extract-N-Amp Plant PCR protocol.^16^ All the direct PCR methods (Method 1, 2, and 3) including Extract-N-Amp are much simpler than DNeasy Powersoil HTP 96 kit which is a 32 step process as described in the manufacturer’s protocol. Accordingly, the total time needed to perform any of the direct PCR methods is much shorter than Powersoil. Method 1 requires negligible time. Method 2 and 3 take 45 minutes and 2 hours, respectively. Extract-N-Amp takes 25 minutes. And Powersoil takes at least 4 hours for extracting gDNA from a 96 well plate. Although Method 3 takes a total of 2 hours, 1 h 45 min are occupied by thermal cycling. The hands-on time only takes 15 min. Method 3 is 10 times simpler (achieving 3 steps and 15 min hands-on time for 96 samples) than Powersoil (32 steps and 3.5 h hands-on time for 96 samples). In addition, since the operation in Method 3 is either reagent (IGEPAL or proteinase K) addition or thermal cycling, all the steps are compatible with automation.

## Discussion

The accuracy and efficiency of the direct PCR methods increases from Method 1 to Method 3. In Method 3 (IGEPAL + Freeze - thaw + Proteinase K), IGEPAL and proteinase K disrupt the phospholipids and membrane proteins; freeze-thaw and heating mechanically disrupt the bacterial membrane. The efficiency of Method 3 is comparable to the widely used methods - DNeasy Blood and Tissue DNA extraction kit and DNeasy Powersoil DNA extraction kit. Particularly, Method 3 exhibits higher efficiency than DNeasy Blood and Tissue DNA extraction kit on disrupting gram negative bacteria - *Escherichia coli* and *Pseudomonas putida*. Importantly, Method 3 (IGEPAL + Freeze - thaw + Proteinase K) is 1600 times cheaper and 10 times faster to execute (in terms of hand-on time) than DNeasy Powersoil DNA extraction kit. The direct PCR methods are also compatible with automation and miniaturization. Because Method 3 only contains IGEPAL and proteinase K addition, freeze-thaw, and thermal cycling, the protocol could be easily integrated into automated laboratory robots. And, because high throughput cultivation assay of microbial communities tend to be performed using smaller volumes, such as in droplets or in 1536 well plates, current column based or beads based DNA extraction methods will be incapable of processing these kinds of samples due to the significant DNA loss for tiny volume and low cell density samples.^7–10, 24^ The direct PCR method will solve this problem by keeping all the gDNA in the PCR without any loss. While the efficiency and precision are the same, the choice of methods are guided largely by the cost, time, and potential for automation.^25^ Thus the direct PCR method is a promising way of microbial community analysis.

Achieving a perfect quantification of a bacterial community to unbiasedly reflect the true composition is extremely difficult, even for the most popular and conventional Powersoil. These reasons include: different bacteria releasing gDNA at different time scale (due to the complexity of bacterial structures) causing the gDNA that is released early in the process to be at risk of being degraded; certain bacteria types being difficult to lyse; and bias via PCR amplification, sequencing, and bioinformatics.^5^ Unsurprisingly, the direct PCR Method 3 has bias too. The bacterial disruption bias of the direct PCR method 3 is likely due to the lack of specific peptidoglycan disruption reagents: such as lysozyme. Because mechanical disruption such as freeze-thaw could theoretically disrupt peptidoglycan, Method 3 only has difficulties to disrupt bacteria with very thick peptidoglycan, such as late stationary phase gram positive bacteria, which is consistent to our results - the qPCR results demonstrate low efficiency of the direct PCR methods on disrupting stationary phase *Lactocuccus lactis* and *Lactobacillius brevis*, but similar efficiency on disrupting exponential phase *Lactocuccus lactis* and *Lactobacillius brevis* compared with DNeasy Blood and Tissue DNA extraction kit. To improve the disruption of peptidoglycan, an enzyme treatment followed by the proteinase K denaturation of this enzyme is a promising solution. Although denatured lysozyme has been shown to inhibit PCR, figuring out the reason of the inhibition would help with designing lysozyme treatment in the direct PCR. In addition, looking for other enzymes that don’t influence PCR would be another way to improve the direct PCR method. In addition, since the quantification bias could be caused not only by DNA extraction, but also by PCR amplification, sequencing, and bioinformatics,^5^ it is important to systematically improve all the steps to better estimate absolute quantification.

The existence of PCR inhibitors in various samples is the limitation of this direct PCR method. Since the chemical composition of microbiome samples is extremely diverse, it is impossible to have a general protocol fitting all the samples. Additional treatment may be required based on our direct PCR method 3. For example, to process the samples with low pH, high ion samples, or soluble PCR inhibitor such as glycerol, it is necessary to remove the liquid phase by centrifuging to pellet the sample. However, in the case of the rich and complex soil samples with high humic acid and unknown PCR inhibitors, DNA extraction-based methods such as Powersoil are recommended.

One of the potential applications of the direct PCR method is to study the bacterial interaction and community assembly. Because the recent advances in sequencing-based quantification have extended its ability from only obtaining relative bacterial abundance to quantify the absolute bacterial abundance by adding reference gene,^26, 27^ measuring total gene load,^28^ or using new algorithms,^29^ it provides a unique way to quantify the bacterial dynamics and assembling in complex bacterial communities using genetic markers such as 16S genes.^30^ However, it is still technically very challenging to understanding bacterial interaction and assembling in complex microbial communities, since the sample number increases dramatically as the increase of the number of species, time point, replicates, and initial conditions. To understand the interaction and assembly of a 7-member community, it requires 127 samples with all the combinations of each species for just one time point without replicates. The number goes to 255 for a 8-member community, and will explode by adding more time points, more replicates, and various initial ratios of each cultivation. This low-cost, automation friendly direct PCR assay is able to handle increasingly large number of microbial community samples, allowing us to study the dynamics, microbial interaction and assembly in a high throughput fashion.

Our direct PCR methods have been evaluated by lab microbial enrichment, microbial community standard culture, and environmental water samples. Since there are diverse microbiome samples with their unique chemical composition, we are not able to screen all types of microbiome samples. Thus, we recommend a PCR/qPCR evaluation to check the efficiency of the direct PCR method on particular types of samples we did not cover. For example, our direct PCR method (Method 3) is shown to successfully extract gDNA from 10 day *E. coli* biofilm samples (Figure S4).

In summary, we provide a cost-effective, simple, and automation friendly direct PCR method for high throughput microbial community composition analysis. This successfully demonstrates the possibility of using direct PCR methods on microbial community analysis. The direct PCR Method 3 (IGEPAL+Freeze-thaw+ProteinaseK) which is comparable to the widely used commercial kits in low cost and quick to do could dramatically increase the throughput of recent 16S sequencing profiling, as well as provide an alternative way to study the microbial community assembly and interactions.

## Methods

### Strains and cultivation conditions

The cell lysis efficiency of direct PCR methods was compared to the Qiagen DNeasy Blood and Tissue kit using the following strains in both stationary phase and exponential phase: *Escherichia coli str. K-12 substr.* MG1655 (*E. coli*), *Pseudomonas putida* KT2440 (*P. putida*), *Lactococcus lactis subsp. cremoris str.* MG1363 (*L lactis*), and *Lactobacillus brevis* ATCC 14869 (*L. brevis*). For the stationary phase cultures, *E. coli* and *P. putida* were inoculated from glycerol stock into LB Lennox medium and cultivated at 37°C with 200 rpm overnight. *L. lactis* and *L. brevis* were inoculated from glycerol stock in MRS medium and cultivated at 30°C without shaking. The OD_600_s of the overnight cultures of *E. coli*, *P. putida*, *L. lactis*, and *L. brevis* were 2.33, 2.28, 1.77, and 3.68, respectively. For exponential phase cultures, overnight cultures were diluted (1:100) from overnight cultures. Culture conditions for all the strains remained the same. Cultures were collected between 4 and 6 hours later. The OD_600_s of exponential phase cultures of *E. coli*, *P. putida*, *L. lactis*, and *L. brevis* were 0.86, 0.75, 0.39, 0.16, respectively. Pellets were stored at −80°C awaiting genomic DNA extractions.

### Mock microbial community standard

ZymoBIOMICS microbial community standards (Zymo Research) were used in this study to quantitatively evaluate the efficiency of direct PCR methods and DNeasy Powersoil kit. This microbial community standard is a defined composition comprised of 5 gram positive bacteria (*Listeria monocytogenes*, *Bacillus subtilis*, *Lactobacillus fermentum*, *Enterococcus faecalis*, *Staphylococcus aureus*), 3 gram negative bacteria (*Pseudomonas aeruginosa*, *Escherichia coli*, *Salmonella enterica*), and 2 yeast strains (*Saccharomyces cerevisiae*, and *Cryptococcus neoformans*). The ZymoBIOMICS microbial community standards were stored at −80°C until use. To avoid the effects of glycerol on microbial community standards cell lysis, 40 ul of the standard was thawed and centrifuged at 10000 g for 5 min. The supernatant was discarded and the pellet was resuspended by 80 ul Milliq/DNase free water immediately before direct PCR or gDNA extraction.

### Environmental groundwater samples

Environmental groundwater samples (GW822D, GW823E) were collected in April 2019 from two uncontaminated background area wells in Bear Creek Valley watershed of Oak Ridge, TN. Groundwater (50ml) was vacuum filtered over 0.2uM pore size and the concentrated microbial community was resuspended in 10 ml PBS. This suspension was used for direct PCR or genomic DNA extraction by DNeasy Powersoil kit.

### Direct PCR methods

We tested three variations of direct PCR methods using the detergent IGEPAL as the core lysis reagent. In the IGEPAL only method (Method 1), IGEPAL detergent was added to PCR (0.1% final concentration) with an initial activation set at 98oC for 10 min, the same as many hot start DNA polymerases. In the IGEPAL+Freeze-thaw method (Method 2), PCR template was prepared by adding an equal volume of bacterial cell suspension to IGEPAL(0.5% final concentration) in a PCR plate, samples were mixed ten times by pipetting up and down, followed by five cycles of freeze-thaw by placing the plates in the −80°C for 15 minutes and then allowing them to thaw at room temperature for 15 minutes. In the IGEPAL+Freeze-thaw+ProteinaseK method (Method 3), samples are prepared in the same way as in the IGEPAL+Freeze-thaw method, except proteinase K (20 mg/mL, Qiagen) was added to the samples after the freeze-thaw cycles, in 96-well PCR plate to reach the final concentration 100 ug/mL. The plates were sealed and centrifuged at 300 x g for 1 minute to collect the liquid to the bottom of each well. Then the samples were treated at 60oC for 1 h and then enzyme deactivated 95oC for 15 min in a thermocycler (Bio-rad). After the treatment, the sample is ready to use as a DNA template in following PCR.

### Genomic DNA extraction methods

Two conventional genomic DNA extraction methods were used in this study for comparison with the direct PCR methods: DNeasy blood and tissue kit (Qiagen) and DNeasy Powersoil kits (Qiagen). DNeasy blood and tissue kit was used to extract genomic DNA from the four model strains per the manufacturer’s specifications, with the additional lysozyme treatment for gram positive strains. For the gDNA extraction, 1 ml culture volume was used for the exponential phase cultures and 400 ul of culture volume was used for stationary phase cultures. Samples were eluted in 400 ul water and stored at −20°C. The bacterial cultures used for gDNA extraction were aliquoted and stored at −80°C. We used DNeasy Powersoil kits to extract genomic DNA from ZymoBIOMICS microbial community standards and microbes from groundwaters as per the manufacturer’s specifications. Three replicates were applied to each sample.

### Quantitative PCR (qPCR)

Quantitative PCR (qPCR) was used for comparing the efficiency of the direct PCR methods with DNeasy Blood and Tissue kit. The extracted gDNA was from the identical sample which was used for direct PCR method. The gDNA was extracted by the DNeasy blood and tissue kit from 400 ul stationary phase culture and eluted by 400 ul elution buffer, so 1 ul extracted gDNA was equivalent to 1 ul original stationary phase culture. For the exponential phase cultures, because the gDNA was extracted from 1 ml cultures and eluted by 400 ul elution buffer, 1 ul extracted gDNA was equivalent to 2.5 ul original exponential phase culture (1 ml cultures volume: 400 ul elution volume = 2.5). The extracted gDNA concentrations were quantified by the Quant-iT double-stranded DNA (dsDNA) assay kit. Each qCPR was conducted in a 20 ul reaction with 10 ul Sso Advanced Universal SYBR Green Supermix (2X), 1.5 ul primer 534F (5 mM), 1.5 ul primer 783R (5 mM), 0.4 ul DNase A (100 mg/ml, Qiagen), 2 ul 1% IGEPAL CA-630, and either 1 ul extracted gDNA or equivalent cell cultures pretreated by the direct PCR methods. Each condition was replicated at least three times. The qPCR was performed using a Bio-rad CFX96 real-time PCR machine with denaturation at 98°C for 10 min, followed by 38 cycles of denaturation at 98°C for 30s and annealing/elongation at 60°C for 1 min 30 s. The melting curve was tested from 60°C to 95°C with 0.5°C/cycle with increments of 5 s per cycle. The Ct was calculated by the Linear Regression method (Lin-Reg).

### 16S amplicon PCR and sequencing

The community structure of the ZymoBIOMICS microbial community standard and two environmental groundwater samples were measured using 16S V3-V4 region Illumina amplicon sequencing, using DNeasy Powersoil - a widely used traditional DNA extraction method, and the direct PCR methods. Primers used in the 16S amplicon PCR were constructed with TruSeq Illumina adapters, barcodes, phasing, and linker sequences, with 341F and 806R targeting 16S V3-V4 hyper variable region of the 16S gene, adopted from Justice et. al.^31^ Genomic DNA extracted from the Powersoil kit was quantified by the Quant-iT dsDNA assay kit. The gDNA of ZymoBIOMICS microbial community standards was between 5.4 ng/ul and 6.0 ng/ul. And the gDNA of water sample GW822D and GW823E was between 0.31 ng/ul and 0.38 ng/ul. The template for the 16S amplicon reaction was either the extracted gDNA from Powersoil kit or direct PCR method treated cells. KAPA HiFi Hotstart ReadyMix was used in the PCR. The PCR was conducted in an 30 ul reaction with 15 ul KAPA HiFi HotStart ReadyMix (2X), 3 ul 1% IGEPAL CA-630, 0.6 ul RNase A (100 mg/ml, Qiagen), 3 ul forward and reverse primer mix (2.5 uM for each), and 8.4 ul template (either extracted gDNA or equivalent cells treated by direct PCR methods) [Note: Specifically, we recommend to use KAPA HiFi Hotstart polymerase, since we found either RNase A or denatured RNase A inhibits PCR by invitrogen Plantinum polymerase but not KAPA HiFi Hotstart polymerase. If you chose modified protocol without RNase A, it is fine to use invitrogen Plantinum polymerase.]. The cycling conditions were 98°C for 10 min, followed by 26 cycles of 98°C for 20 s, 53°C for 30 s, and 72°C for 2 min, and a final extension step of 72°C for 5 min. The PCR products were run on 1.2 % agarose gel to determine the amplicon concentration. Based on the quantification from gel imaging, similar amounts of PCR products of each PCR were pooled together and purified with AMPure XP beads per manufacturer’s protocol and the purified product was quantified by the Quant-iT dsDNA high sensitivity assay. Then 4 nM library stock was prepared by diluting the purified amplicon DNA with water. The library stock was denatured and diluted following manufacturer’s instruction, a final concentration of 20 pM denatured library was loaded onto the flowcell and sequenced using a custom read 2 primer (5’- CGGTCTCGGCATTCCTGCTGAACCGCTCTTCCGATCT). Amplicons were sequenced using Illumina 600 bp v3 kit with 350 bp read 1 and 250 bp read 2 on the Illumina Miseq platform.

### Sequencing data processing and analysis

All the analysis are performed in R (3.6.0). The amplicon sequence data was analyzed using DADA2.^32^ Most forward and reverse reads did not meet the criteria for high quality assembly during the reads merging (less than 1 mismatch and 20 bp overlap-find actual parameter used for assemble) due to the low quality sequence of the overlapping region in read 2, so the DADA2 was analyzed by read 1 only, which includes V3 and V4 region. Primer sequences were trimmed with cutadapt, sequence length was trimmed to 300 bp, low quality reads were filtered by the settings (maxN=0, maxEE=3, truncQ=2), and chimeric reads were removed, and the relative abundance of exact sequence variants (ESVs) was calculated by DADA2. The taxonomy was assigned using the naive Bayesian classifier to assign taxonomy across multiple ranks with the SILVA Database V132 as a reference.^33^ ZymoBIOMICS microbial community standard strains were counted by the reads of each species. For the water samples, alpha diversity was calculated based on ESVs using Shannon index. Bray-Curtis distances was calculated using the normalized and log transformed ESV metrics, and examined with PERMANOVA by using the Adonis function with 1,000 permutations both in vegan (2.5-7). And PCoA based on Bray-Curtis distances was used to visualize and compare the different water samples and different methods.

## Acknowledgements

We thank Dan Williams, Kenneth Lowe, Terry Hazen, Dominique Joyner, Katie Walker, Andrew Putt, Regina Wilpiszeski, and Emma Dixon at Oak Ridge National Laboratory for the ENIGMA Spring Sampling to provide water samples GW822D and GW823E. And we thank Yolanda Huang and Sean Carim for assembly of our strain standards and helpful discussion. This work conducted by ENIGMA-Ecosystems and Networks Integrated with Genes and Molecular Assemblies (https://enigma.lbl.gov/), a Scientific Focus Area Program at Lawrence Berkeley National Laboratory, was supported by the Office of Science, Office of Biological and Environmental Research, of the U. S. Department of Energy under Contract No. DE-AC02-05CH1123.

## Author contributions

FS conceived the project and wrote the original draft. FS and AC conducted the experiments. FS and JVK designed the computational framework and analyzed the data. APA supervised the project. FS wrote the manuscript. All authors reviewed and revised the manuscript.

## Additional information

The authors declare no competing financial interests.

## Data availability

The composition of the microbial community standard samples and water samples are provided in Table S2 and Figure S2. The raw sequencing data is made available at Figshare (https://figshare.com/s/d31777db4915ee7563da). The code is available on Github (https://github.com/fasong/Direct_PCR_4_bacterial_community_analysis.git)

## Supplementary Tables

**Table S1.** Average ΔCt and *p* value (*t*-test) from quantitative PCR between gDNA extracted by DNeasy blood and tissue kit and cells treated by direct PCR methods. ΔCt = Ct of direct PCR method - gDNA control extracted by DNeasy blood and tissue kit. (Notes: IGEPAL only is Method 1; IGEPAL + Freeze-thaw is Method 2; IGEPAL + Freeze-thaw + proteinase K is Method 3.)

|  |  | Escherichia Exponential | Escherichia Stationary | Pseudomonas Exponential | Pseudomonas Stationary | Lactococcus Exponential | Lactococcus Stationary | Lactobacillus Exponential | Lactobacillus Stationary |
| --- | --- | --- | --- | --- | --- | --- | --- | --- | --- |
| IGEPAL only | P value | 0.3985 | 0.4587 | 0.1215 | 0.8627 | <0.0001 | <0.0001 | <0.0001 | <0.0001 |
| | $\Delta Ct$ | -1.733 ± 1.835 | 0.4938 ± 0.6028 | -3.568 ± 1.820 | 0.06502 ± 0.3526 | 6.272 ± 1.127 | 2.409 ± 0.4388 | 3.931 ± 0.1825 | 30.18 ± 0.4252 |
| IGEPAL+Freeze-thaw | P value | 0.1918 | 0.1602 | 0.9065 | 0.4247 | 0.0157 | <0.0001 | <0.0001 | <0.0001 |
| | $\Delta Ct$ | 2.147 ± 1.368 | 0.4815 ± 0.3111 | 0.1567 ± 1.253 | -0.4500 ± 0.5067 | 2.562 ± 0.8627 | 3.244 ± 0.4509 | 3.113 ± 0.2443 | 26.87 ± 0.5764 |
| IGEPAL+Freeze-thaw+ProteinaseK | P value | <0.0001 | 0.0012 | 0.0014 | <0.0001 | 0.787 | <0.0001 | 0.1934 | <0.0001 |
| | $\Delta Ct$ | -3.149 ± 0.2015 | -1.748 ± 0.2626 | -4.386 ± 0.6922 | -2.178 ± 0.1866 | 0.1947 ± 0.6993 | 7.884 ± 0.5280 | 0.7051 ± 0.5016 | 12.86 ± 0.3844 |

**Table S2.** Comparison of the relative abundance of ZymoBIOMICS microbial community standards (Control) and those obtained by DNeasy PowerSoil kits and direct PCR methods. (Notes: IGEPAL only is Method 1; IGEPAL + Freeze-thaw is Method 2; IGEPAL + Freeze-thaw + proteinase K is Method 3.)

| Relative abundance (%) | Lactobacillus fermentum | Enterococcus faecalis | Escherichia coli | Salmonella enterica | Pseudomonas aeruginosa | Staphylococcus aureus | Bacillus subtilis | Listeria monocytogenes |
| --- | --- | --- | --- | --- | --- | --- | --- | --- |
| Control | 18.4 | 9.9 | 10.1 | 10.4 | 4.2 | 15.5 | 17.4 | 14.1 |
| PowerSoil_1 | 17.89 | 12.6 | 21.54 | 25.51 | 8.94 | 6.91 | 4.27 | 2.34 |
| PowerSoil_2 | 15.42 | 16.52 | 17.82 | 13.41 | 10.01 | 14.31 | 8.21 | 4.3 |
| PowerSoil_3 | 14.99 | 15.68 | 17.68 | 14.29 | 9.29 | 13.59 | 8.99 | 5.49 |
| PowerSoil_4 | 21.02 | 9.81 | 21.92 | 21.62 | 9.01 | 7.21 | 6.81 | 2.6 |
| PowerSoil_5 | 13.94 | 11.09 | 23.8 | 24.21 | 9.87 | 7.43 | 7.12 | 2.54 |
| PowerSoil_6 | 11.22 | 15.13 | 19.54 | 16.63 | 10.82 | 12.32 | 9.62 | 4.71 |
| IGEPAL only_1 | 67.9 | 14.1 | 5.3 | 2.1 | 1 | 6 | 1.4 | 2.2 |
| IGEPAL only_2 | 60.32 | 16.93 | 5.91 | 2.81 | 1.7 | 8.42 | 1.9 | 2 |
| IGEPAL only_3 | 65.53 | 15.13 | 5.31 | 2.51 | 1.3 | 6.81 | 1.7 | 1.7 |
| IGEPAL+Freeze-thaw_1 | 55.26 | 17.82 | 6.81 | 3.4 | 1.8 | 9.71 | 2.8 | 2.4 |
| IGEPAL+Freeze-thaw_2 | 51.9 | 20.04 | 7.92 | 3.81 | 2 | 9.62 | 2.2 | 2.51 |
| IGEPAL+Freeze-thaw_3 | 52.05 | 20.22 | 7.61 | 3.7 | 1.9 | 9.71 | 2.3 | 2.5 |
| IGEPAL+Freeze-thaw+ProteinaseK_1 | 11.72 | 32.67 | 17.84 | 9.52 | 4.01 | 13.73 | 4.81 | 5.71 |
| IGEPAL+Freeze-thaw+ProteinaseK_2 | 13.3 | 33.5 | 18.5 | 9.9 | 3.3 | 11.9 | 4.4 | 5.2 |
| IGEPAL+Freeze-thaw+ProteinaseK_3 | 16.02 | 32.73 | 16.02 | 9.01 | 2.9 | 12.21 | 5.41 | 5.71 |

**Table S3.**
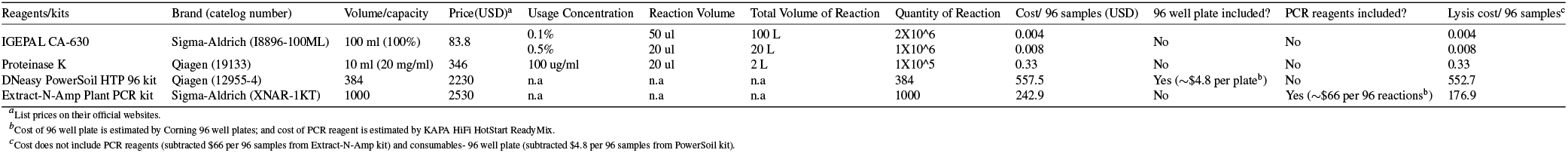
Cost analysis of the lysis reagents of direct PCR methods, DNeasy Powersoil kit, and Extract-N-Amp Plant PCR kit. (Notes: IGEPAL only is Method 1; IGEPAL + Freeze-thaw is Method 2; IGEPAL + Freeze-thaw + proteinase K is Method 3.)

## Supplementary Figures

**Figure S1.**
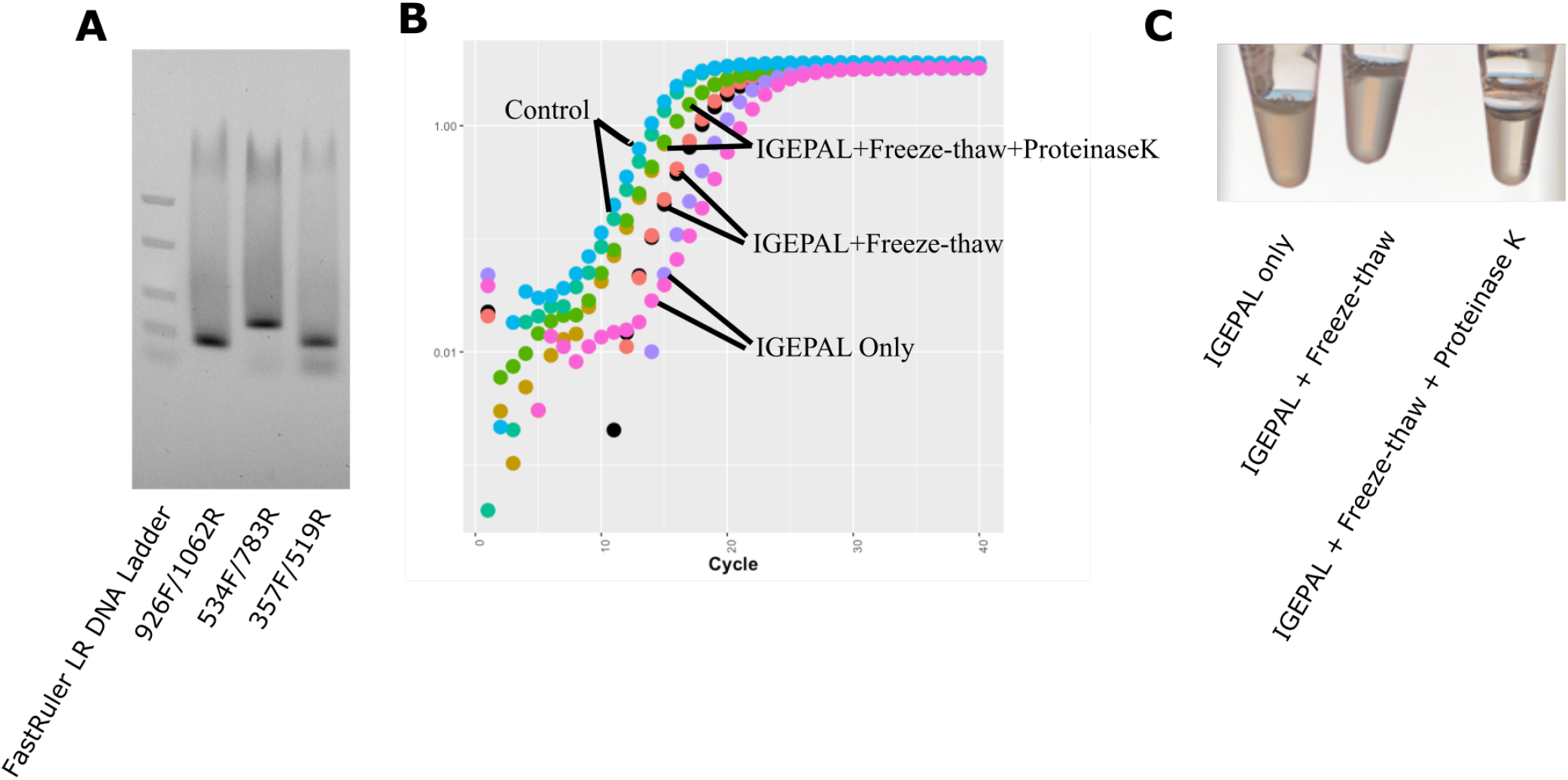
Evaluation of cell lysis efficiency of direct PCR methods using quantitative PCR (qPCR). (A) qPCR products gel images of direct PCR method 3 treated *E. coli* cells using primers 926F/1062R, 534F/783R, and 357F/519R. (B) Amplification curve of qPCR using DNeasy blood and tissue kits extracted gDNA (Control) and equivalent *L lasctis* cells treated by direct PCR methods. (C) The changes of opacity of the cell lysates by direct PCR method 1 (IGEPAL only), method 2 (IGEPAL + Freeze-thaw), and method 3 (IGEPAL + Freeze-thaw + Proteinase K).

**Figure S2.**
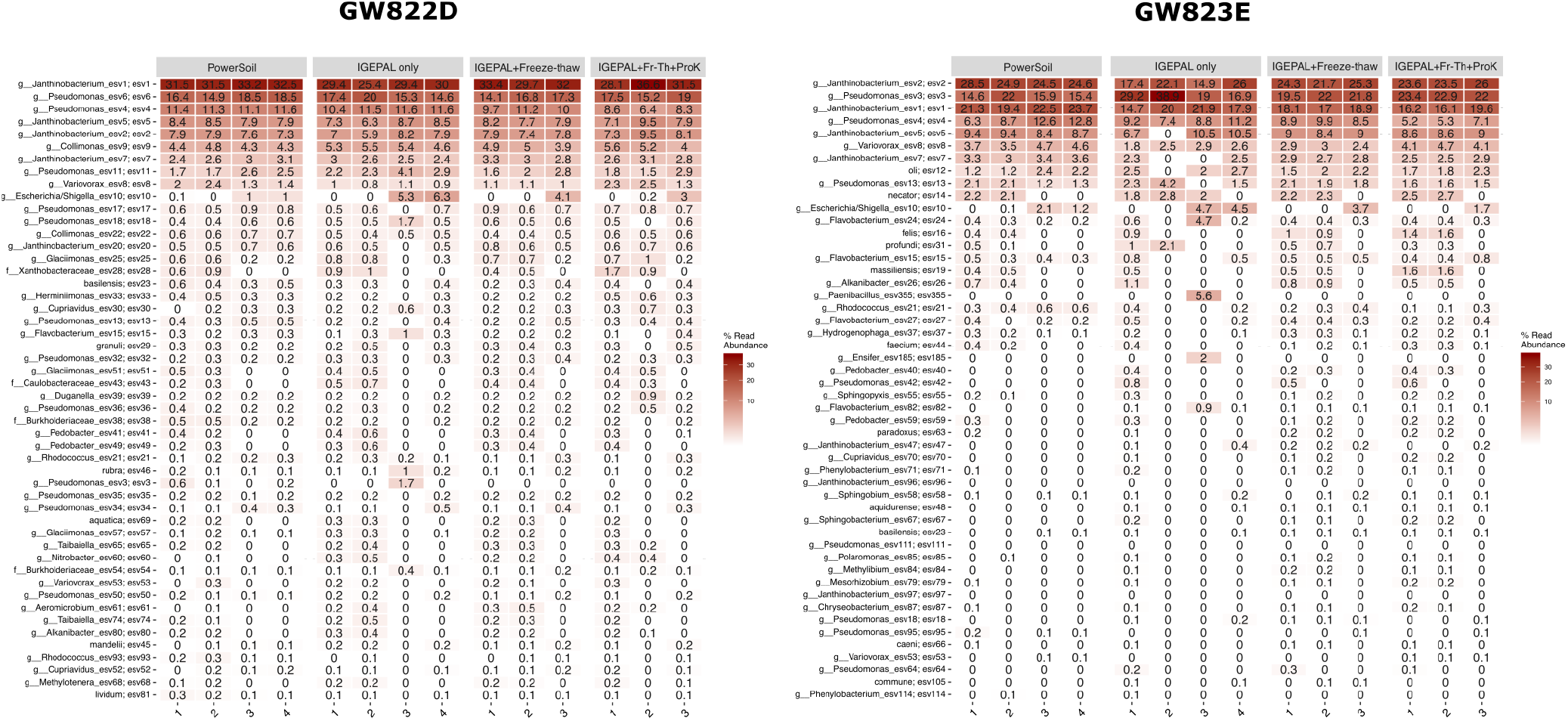
Relative abundance of ESV in ground water samples GW822D and GW823E obtained by DNeasy Powersoil kit and direct PCR methods. (Notes: IGEPAL only is Method 1; IGEPAL + Freeze-thaw is Method 2; IGEPAL + Freeze-thaw + proteinase K is Method 3.)

**Figure S3.**
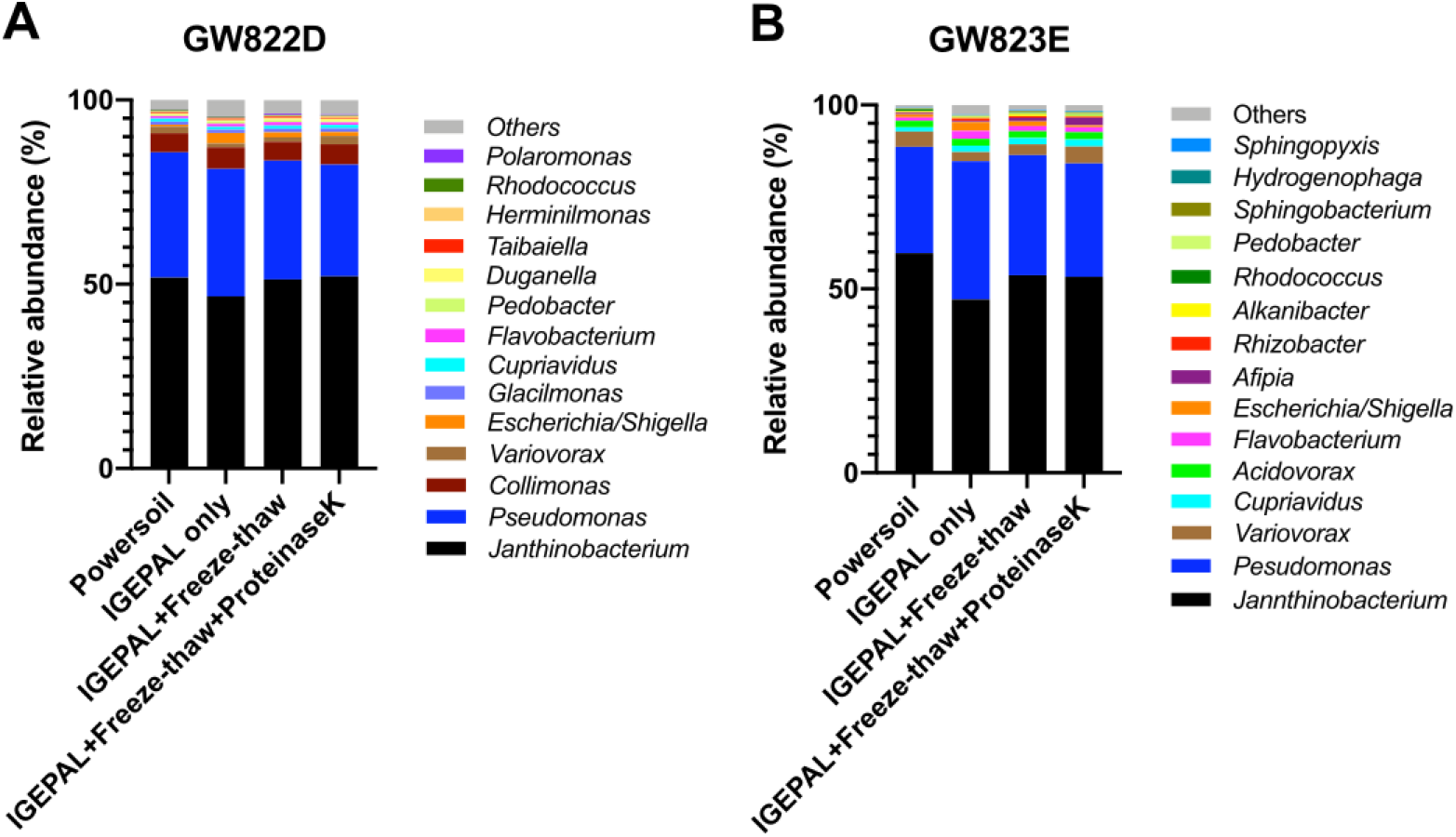
The composition of ground waters obtained by direct PCR methods and DNeasy Powersoil kit. (Notes: IGEPAL only is Method 1; IGEPAL + Freeze-thaw is Method 2; IGEPAL + Freeze-thaw + proteinase K is Method 3.) (A) Average bacterial taxonomic composition of the ground water sample GW822D. (B) Average bacterial taxonomic composition of the ground water sample GW823E.

**Figure S4.**
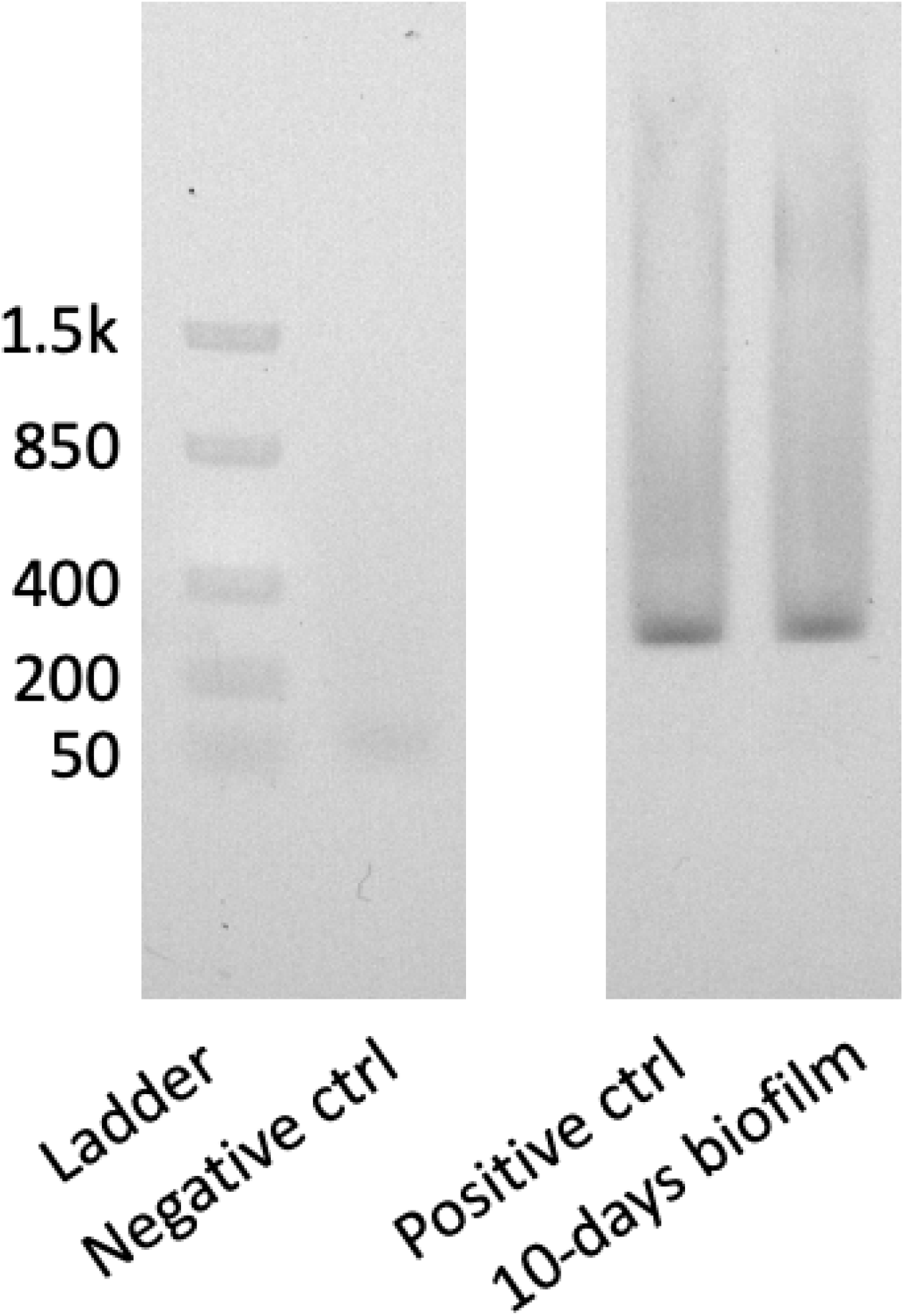
The gel images of the PCR products of 10-days *E. coli* bioflim on LB agar by direct PCR method 3. The PCR was conducted in an 25 ul reaction with 15 ul KAPA HiFi HotStart ReadyMix (2X), 2.5 ul 1% IGEPAL CA-630, 0.6 ul RNase A (100 mg/ml, Qiagen), 2.5 ul forward (534F) and reverse primer (783R) mix (2.5 uM for each), and 2 ul template (a small loop of 10 days *E. coli* biofilms on agar dissolved in 50 ul water and treated by direct PCR method 3 - IGEPAL+Freeze-thaw+ProteinaseK) or 1 ul extracted *E. coli* gDNA (positive control) or water (negative control). The cycling conditions were 95°C for 10 min, followed by 35 cycles of 98°C for 20 s, 55°C for 15 s, and 72°C for 1 min, and a final extension step of 72°C for 10 min.

